# Quantitative Semisolid Magnetization Transfer and Relayed Nuclear Overhauser Effect Imaging in a Multiple Sclerosis Mouse Model Using Deep Magnetic Resonance Fingerprinting

**DOI:** 10.64898/2026.08.17.743476

**Authors:** Ruth Ben Chaim, Michal Rivlin, Or Perlman

## Abstract

Magnetic resonance imaging (MRI) is the imaging modality of choice for the diagnosis, characterization, and monitoring of multiple sclerosis (MS). Nevertheless, the contrasts manifested by MS lesions often overlap with those of other pathological conditions, highlighting the need for additional disease biomarkers. In addition, while saturation transfer (ST) MRI provides molecular information associated with myelin, protein, and lipids, quantifying the underlying proton exchange parameters remains challenging. Here, we describe a strategy that extends and modifies AI-boosted ST magnetic resonance fingerprinting (MRF) imaging at 7T. This approach was used to quantify the dynamics of the semisolid magnetization transfer (MT) and the aliphatic relayed nuclear Overhauser effect (rNOE at −3.5 ppm and −1.6 ppm relative to water) in a longitudinal cuprizone MS mouse model (n=12). In lipid phantoms, the reconstructed proton volume fractions were strongly correlated with known lipid concentrations across all three proton pools (r>0.96, p<0.001). In vivo, semisolid MT and rNOE proton volume fractions in the corpus callosum demonstrated a significant decrease (p<0.01) as early as week 4 of cuprizone feeding, preceding changes detected by conventional water relaxometry. ST-MRF based biomarkers were in agreement with histological findings. Overall, our results demonstrate the feasibility of rapid, multi-pool ST-MRF quantification for MS characterization.

## Introduction

Multiple sclerosis (MS) is a neurodegenerative disease in which an autoimmune response targets the lipid-rich myelin sheath surrounding neuronal axons, causing damage and loss (demyelination).^1^ The disease affects about 3 million people, predominantly young adults, worldwide and may lead to progressive disability and reduced quality of life. While early diagnosis is crucial for timely intervention and optimal management of symptoms,^2^ the period between first seeking medical attention for disease-related symptoms and receiving an MS diagnosis may reach one to three years.^3^

Magnetic resonance imaging (MRI) is the primary imaging modality used for the diagnosis and monitoring of MS. However, despite the use of multiple contrast modalities and periodic updates to imaging criteria,^4^ accurate diagnosis remains challenging, and imaging specificity is often limited.^5^

Saturation transfer (ST) MRI is an increasingly investigated method for acquiring molecular information.^6^ The technique employs frequency selective radiofrequency (RF) pulses to characterize proton exchange rates associated with various targets.^7^ In the context of MS, the semisolid magnetization transfer (MT) effect has long been associated with the concentration of large, immobile macromolecules, such as those found in myelin sheaths.^8,9^ Specifically, both preclinical^10^ and clinical studies^11–13^ have detected lower semisolid MT signals in demyelinating tissue. Additional information can be obtained by exploiting the relayed nuclear Overhauser Enhancement (rNOE), where the magnetization of non-exchangeable protons is transferred to neighboring exchangeable protons via cross-relaxation and then relayed to bulk water by chemical exchange.^14^ Mobile macromolecules such as lipids and proteins can provide a detectable rNOE effect that reflects myelination dynamics at a chemical shift of −3.5 ppm relative to water.^15,16^ In addition, a distinct rNOE signal at −1.6 ppm has recently been reported.^17–20^ Although to the best of our knowledge, this signal has not previously been studied in the context of MS, it has been linked to membrane phospholipids, specifically phosphatidylcholines,^20^ which are major structural constituents of myelin membranes.^21^ This suggests that the −1.6 ppm rNOE signal may provide complementary information regarding myelin integrity.

Unfortunately, despite the promise of ST MRI, standard analytic methods, such as magnetization transfer ratio asymmetry (MTR_asym_), are inherently influenced by contributions from multiple proton pools and water T_1_ relaxation, while failing to quantify proton volume fractions and exchange rates. Furthermore, traditional Z-spectrum acquisition is relatively time consuming, requiring measurements across numerous saturation frequency offsets with relatively long saturation and recovery periods.^22^

ST magnetic resonance fingerprinting (MRF) is an emerging approach for accelerating quantitative ST imaging.^23^ This technique utilizes a pseudo-random series of saturation pulses within a non-steady state acquisition protocol to encode different combinations of proton exchange parameters into unique signal trajectories (“fingerprints”).^24^ Acquired signals are then decoded into molecular parameter maps by comparing them to a simulated dictionary generated by a Bloch-McConnell simulator. Training neural networks (NN) to map a smooth representation of the proton exchange parameter manifold, can accelerate the quantification and overcome the limitations of a discrete dictionary.^25,26^ Indeed, ST-MRF was recently leveraged for molecular characterization of metastatic brain tumors^23^, oncolytic virotherapy,^27^ and Parkinson’s Disease^28,29^. These studies highlight the potential of ST-MRF for non-invasive molecular imaging across a range of pathological conditions.

Here, we describe a method that uses deep ST-MRF for simultaneous quantification of the proton exchange dynamics of three proton pools: semisolid MT; rNOE at −3.5 ppm; and (for the first time) rNOE at −1.6 ppm (**Figure 1**). We then apply this framework to characterize the proton exchange dynamics of these pools in a longitudinal MS mouse model. Cuprizone feeding is a well-established model of demyelination that selectively affects myelin, particularly within the corpus callosum.^30^ Importantly, withdrawal of cuprizone results in spontaneous remyelination, providing a unique opportunity to investigate both demyelination and recovery using quantitative ST-MRF biomarkers.

**Figure 1.**
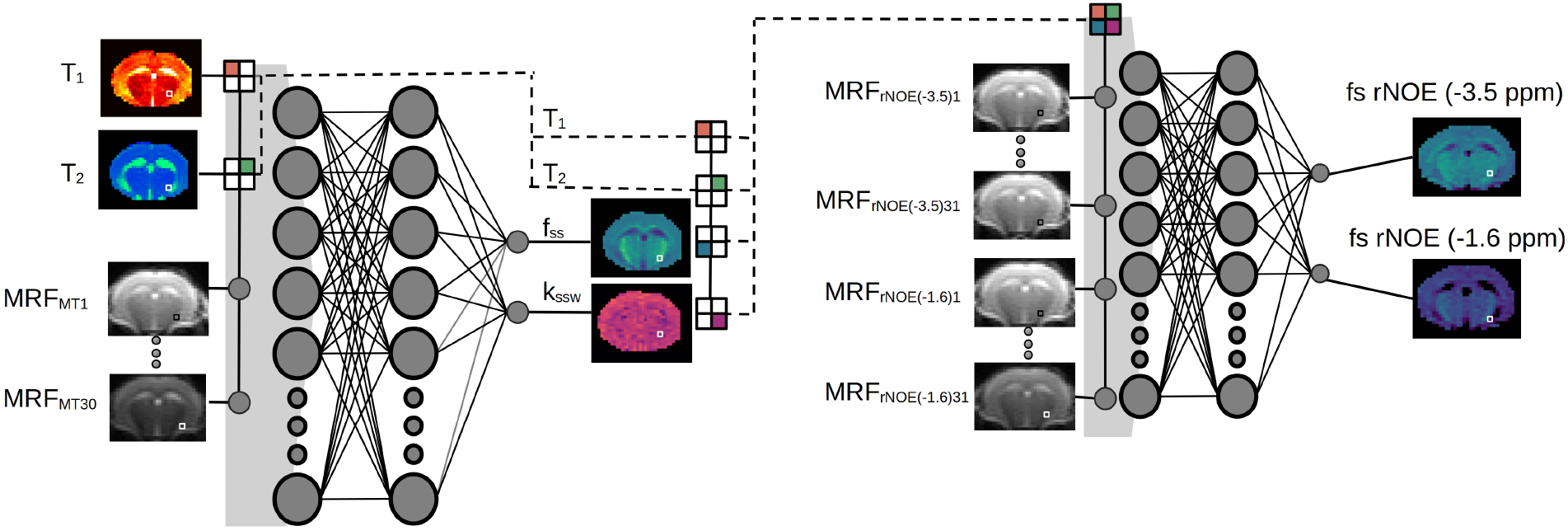
Schematic representation of the serial deep neural network (NN) pipeline. The framework consists of two concatenated steps, each employing a fully connected NN with two 300-node hidden layers. The first NN receives quantitative water T_1_ and T_2_ maps as input along with 30 raw ST-MRF semisolid MT encoding images (in a voxelwise manner). The resulting semisolid MT proton volume fraction (f_ss_) and exchange rate (k_ssw_) are then fed into a second NN, which integrates them with 62 raw rNOE (at both −3.5 ppm and −1.6 ppm) encoding images to provide the respective proton volume fractions (f_s,_).

## Materials and Methods

### Phantom Study

Phantoms were prepared from porcine brain lipid extracts (Avanti Polar Lipids, catalog no. 131101). The extract comprises the major lipid classes of the human brain, including phosphatidylcholine (PC), phosphatidylethanolamine (PE), phosphatidylinositol (PI), phosphatidylserine (PS), phosphatidic acid (PA), and cholesterol. The lipid extract was diluted in phosphate buffered saline (10mM PBS, pH 7.00) and stirred at 65 °C to produce a homogeneous dispersion. Seven 1 ml vials containing lipid concentrations ranging from 3–24% (w/v) were prepared and placed together inside a 50 ml Falcon test tube. The concentration range was selected to approximate the physiological range of myelin lipid content in white matter.^31^ Phantoms were scanned at 37°C, maintained using a hot air blower with temperature feedback control.

### Animal Study

All experimental procedures were approved by the Tel Aviv University Institutional Animal Care and Use Committee (IACUC) and adhered to the Israel National Research Council (NRC) ethical principles. Male C57BL/6J mice (n=12) were acquired from Harlan Israel and were housed in the animal facility of Tel Aviv University under standard conditions. Following a baseline MRI scan, all mice were fed a diet containing 0.3% cuprizone mixed into ground chow. This concentration has been shown to induce demyelination while maintaining animal safety.^32^

After 7 weeks of cuprizone administration, five mice were sacrificed for histological analysis, while one additional mouse was excluded due to an unexpected death unrelated to the cuprizone treatment. The remaining six mice were returned to a standard diet for an additional 5 weeks to allow spontaneous remyelination. At week 12, these mice were sacrificed for histological analysis. An additional group of three mice served as untreated histological controls and were sacrificed at baseline. The complete experimental timeline is presented in **Figure 2**.

**Figure 2.**
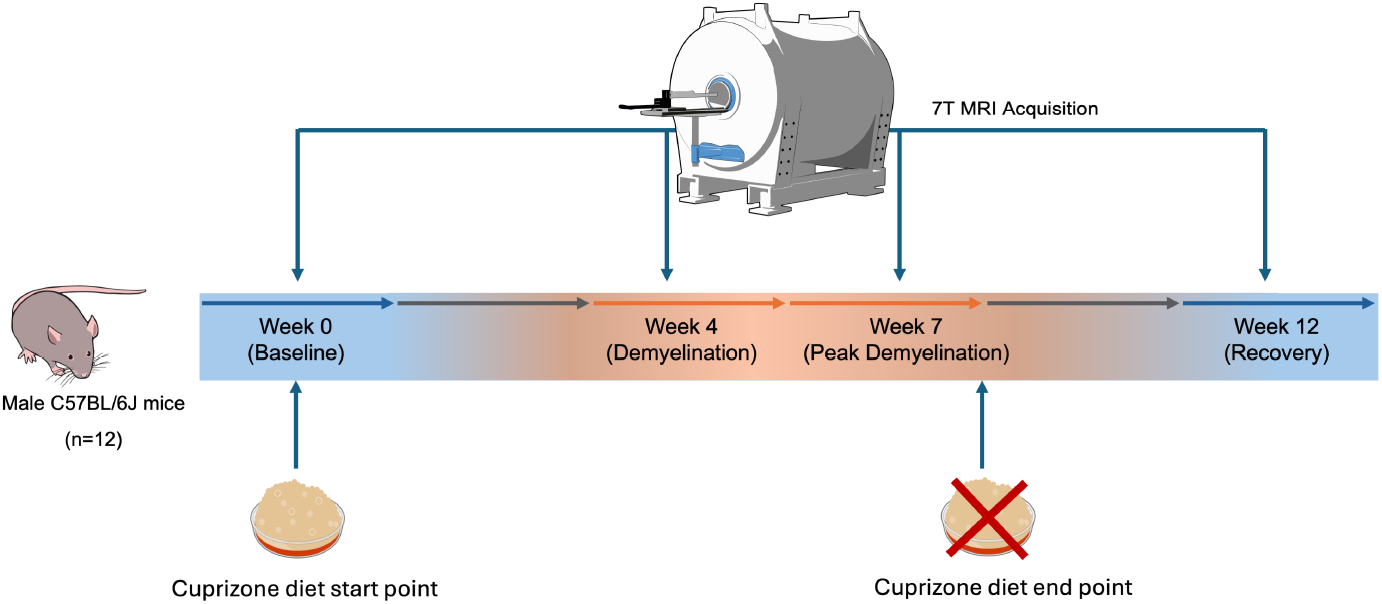
Experimental timeline describing the dietary supply and imaging time points across the 12-week study. Imaging was performed at baseline, week 4 (early demyelination), week 7 (peak demyelination), and week 12 (remyelination).

### MRI Acquisition

Imaging was performed using a preclinical 7T scanner (Bruker, Germany). Three spin echo planar imaging (SE-EPI) based ST-MRF acquisitions^33^ were sequentially applied to encode the semi-solid MT, rNOE (−3.5 ppm), and rNOE (−1.6 ppm) proton pool information into unique signal trajectories. The protocols consisted of 30 or 31 raw images acquired by varying the saturation pulse power between 0-4 μT (**Figure S1**).^34^ Saturation frequency offsets were varied between 6 and 14 ppm for semisolid MT encoding,^27^ with fixed frequency offsets of −3.5 ppm and −1.6 ppm used for rNOE encoding. The saturation pulse time was 2.5 s, the flip angle (FA) was 90°, and the recovery time was 1 s for all MRF protocols, with an additional unsaturated image acquired first (TR = 15 s) for the two rNOE protocols.^28^ The total acquisition time for all three ST-MRF protocols was 5 minutes and 45 seconds. Water T_1_ and T_2_ maps were acquired using rapid acquisition with relaxation enhancement (RARE) and multi slice multi echo (MSME) sequences, respectively. Two full Z-spectra were acquired for comparison with conventional saturation transfer imaging. The first was optimized for rNOE, amide, and semisolid MT contrast, while the second employed an amine and amide concentration-independent detection (AACID)-based acquisition scheme for pH-sensitive imaging.^35^ The saturation pulse powers and durations were 0.7/1.5 μT and 3/4 s, and the acquisition times were 928/1152 s, for the slow-exchange/AACID schedules, respectively. The frequency offset spanned the range from +7 to −7 ppm with step sizes of 0.25 and 0.2 ppm, respectively. These acquisitions employed the same SE-EPI readout module as the ST-MRF protocol (FA = 90°, TE = 20 ms), yet with a TR = 8000 ms, and number of averages (NA) = 2.

### ST-MRF Dictionary Generation

Synthetic ST-MRF signal trajectories were generated using a numerical Bloch-McConnell solver implemented in C++ with a Python front-end.^33^ The simulation yielded a comprehensive dictionary containing a total of 34,935,072 entries within 63 min, using a computing server employing up to 100 CPU workers. This dictionary was used for training the deep learning quantification model. Complete in vivo simulation parameters are available in **Tables S1-S2**.

### AI-Based Parameter Quantification

Deep-learning-based quantification was obtained using a serial reconstruction pipeline^26,27^ composed of two reconstruction networks. The first network was optimized for semi-solid MT quantification, and the second targeted the two rNOE (−3.5 ppm and −1.6 ppm) component quantification, while using the estimated voxelwise semisolid MT parameters to narrow down the parameter space. Each NN comprised two hidden layers with 300×300 neurons each (**Figure 1**), and received the relevant raw MRF data, alongside the water T_1_ and T_2_ values (estimated separately) as input.^27^ The pipeline was implemented in PyTorch and trained on an NVIDIA L40S GPU. Total training time took approximately 10 h, while inference took less than 1 s per mouse dataset.

### Statistical Analysis

The corpus callosum region of interest (ROI) was manually delineated using the Allen mouse Brain Atlas as anatomical reference.^36^ Pearson’s r and associated two-tailed p-values were calculated using the SciPy library for Python. Group comparative analysis used one-way ANOVA, followed by correction for multiple comparisons using a two-sided Tukey’s multiple comparisons test, calculated using the statsmodels library for Python.^37^ Statistical significance was defined as p < 0.05.

### Histological Analysis

Formalin-fixed paraffin-embedded (FFPE) brain tissue sections were cut at a thickness of 5 μm. Sections were then de-paraffinized and immunostained for specific markers. The primary region of interest for all evaluations was the corpus callosum (as defined in the Allen Brain Atlas).^36^ Immunofluorescence used a calibrated antigen retrieval (AR) method at pH 6.0. Double fluorescence immunostaining used Chicken pAb anti-GFAP (#Ab4674, Abcam; at a concentration of 13.19 μm/mL specific IgY) to target astrocytes, and Mouse MBP at a concentration of 20 μm/mL) to target myelin basic protein. Non specific background and negative controls included only the relevant secondary antibody: Donkey anti-Mouse Cy3 or Donkey anti-Chicken 488. Normal, positive staining for the astrocyte marker (GFAP) was used to confirm tissue and antibody integrity. Images were acquired with an Olympus fluorescence scanner at 20x magnification. All images for a given antibody were acquired under identical exposure conditions to enable comparison across all samples and treatment groups.

## Results and Discussion

### ST-MRF in Brain Lipid Phantoms

An initial brain lipid phantom study of seven phantoms containing different concentrations of porcine brain lipid extract was conducted as validation of the proposed ST-MRF simultaneous quantification of three proton volumes: semisolid MT; rNOE at −3.5 ppm; and rNOE at −1.6 ppm. The extract comprises the major lipid classes found in the human brain^38,39^ (see methods section), many of which have been implicated in the pathological processes associated with MS.^40-42^ **Figure 3a** shows the deep MRF reconstructed proton volume fraction maps of the porcine brain lipids, demonstrating a clear proton volume fraction dependency on the total lipid concentration for all proton pools. **Figure 3b** analyzes the correlation between the reconstructed proton volume fractions and the total brain lipid concentration. Pearson’s r values demonstrated a strong positive correlation in all cases: r = 0.969, p < 0.001 for semisolid MT; r = 0.985, p < 0.0001 for rNOE at −3.5 ppm; and r = 0.989, p < 0.0001 for rNOE at −1.6 ppm. These findings are consistent with previous in vitro ST-MRF studies, including BSA-phantoms for rNOE quantification.^26,43^

**Figure 3.**
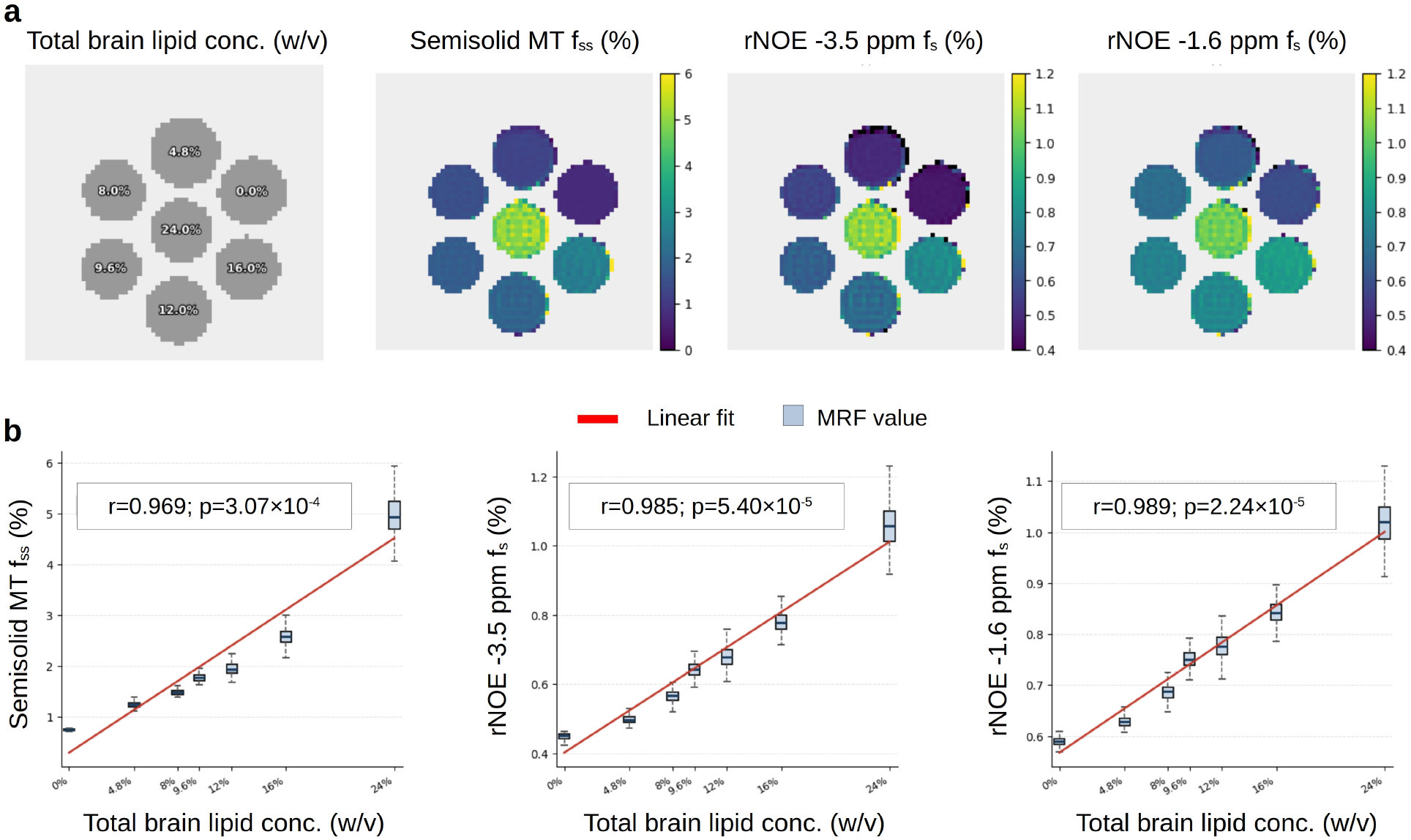
Deep MRF-based proton volume fraction quantification in porcine brain lipid phantoms. **a**. Quantitative proton volume fraction maps (**right**) alongside the corresponding total brain lipid concentrations (**left**). **b**. Statistical analysis of the correlation between the quantitative proton volume fractions and total brain lipid concentration.

### ST MRF in a Cuprizone Mouse Model

Longitudinal deep MRF-based proton volume fraction maps from a representative mouse brain are shown in **Figure 4c-e, f-h**.

**Figure 4.**
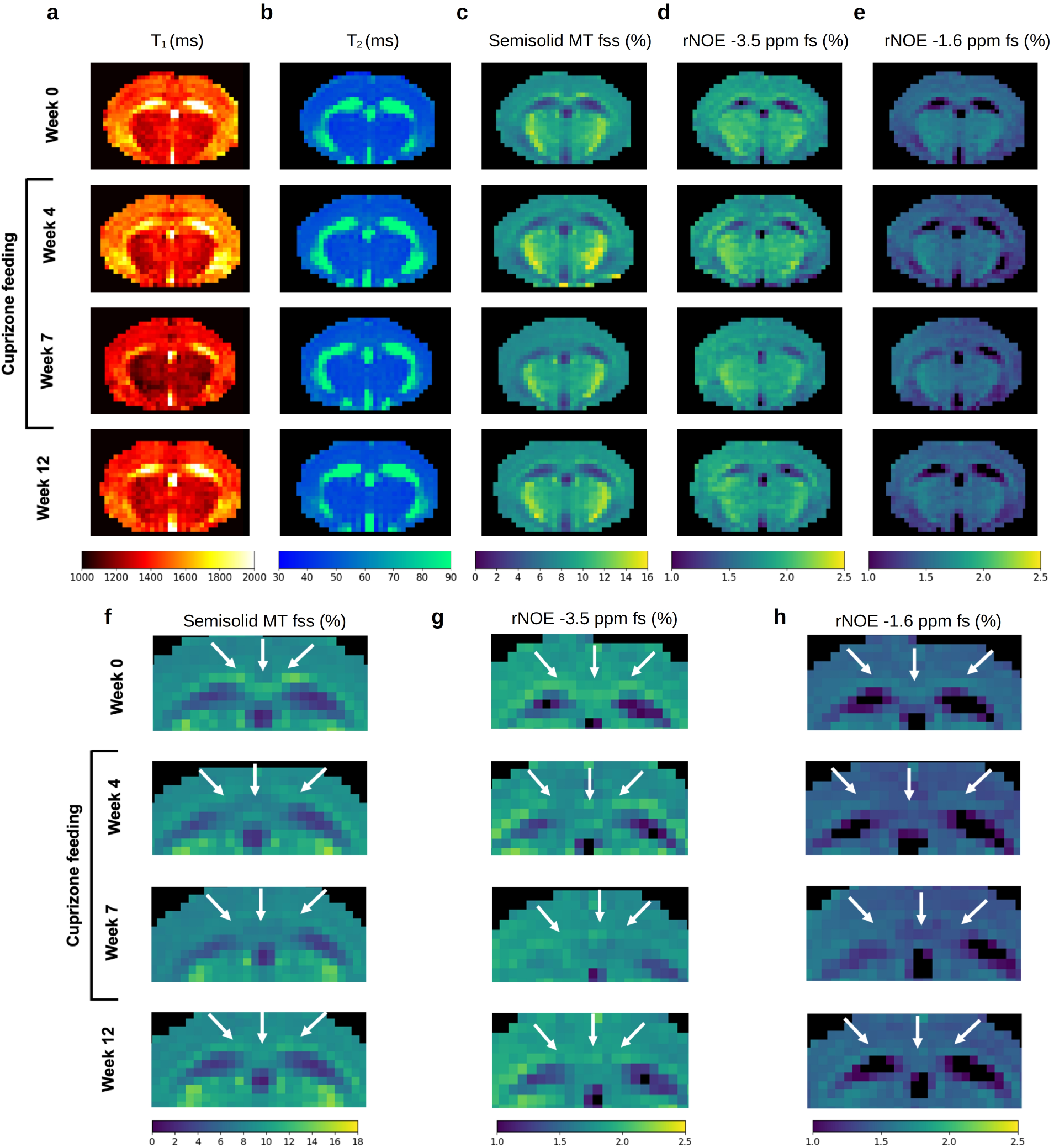
Quantitative parameter maps from a representative cuprizone treated mouse. Each row shows an experimental time point (top to bottom): baseline (week 0), early demyelination (week 4), peak demyelination (week 7), and remyelination (week 12). Water T_1_ (**a**) and T_2_ (**b**) are presented alongside the deep MRF-based proton volume fraction maps of the semisolid MT (**c**), rNOE at −3.5 ppm (**d**), and rNOE at −1.6 ppm (**e**). (**f-h**). A zoom-in view across all time points for the semisolid MT (**f**), rNOE at −3.5 ppm (**g**) and rNOE at −1.6 ppm (**h**). The arrows point to the corpus callosum region.

There was a marked decrease in the proton volume fraction compared to baseline through weeks 4 and 7 in all cases, followed by a trend to recovery at week 12. A quantitative statistical analysis across the entire mouse cohort at all four time points is available in **Figure 5**.

**Figure 5.**
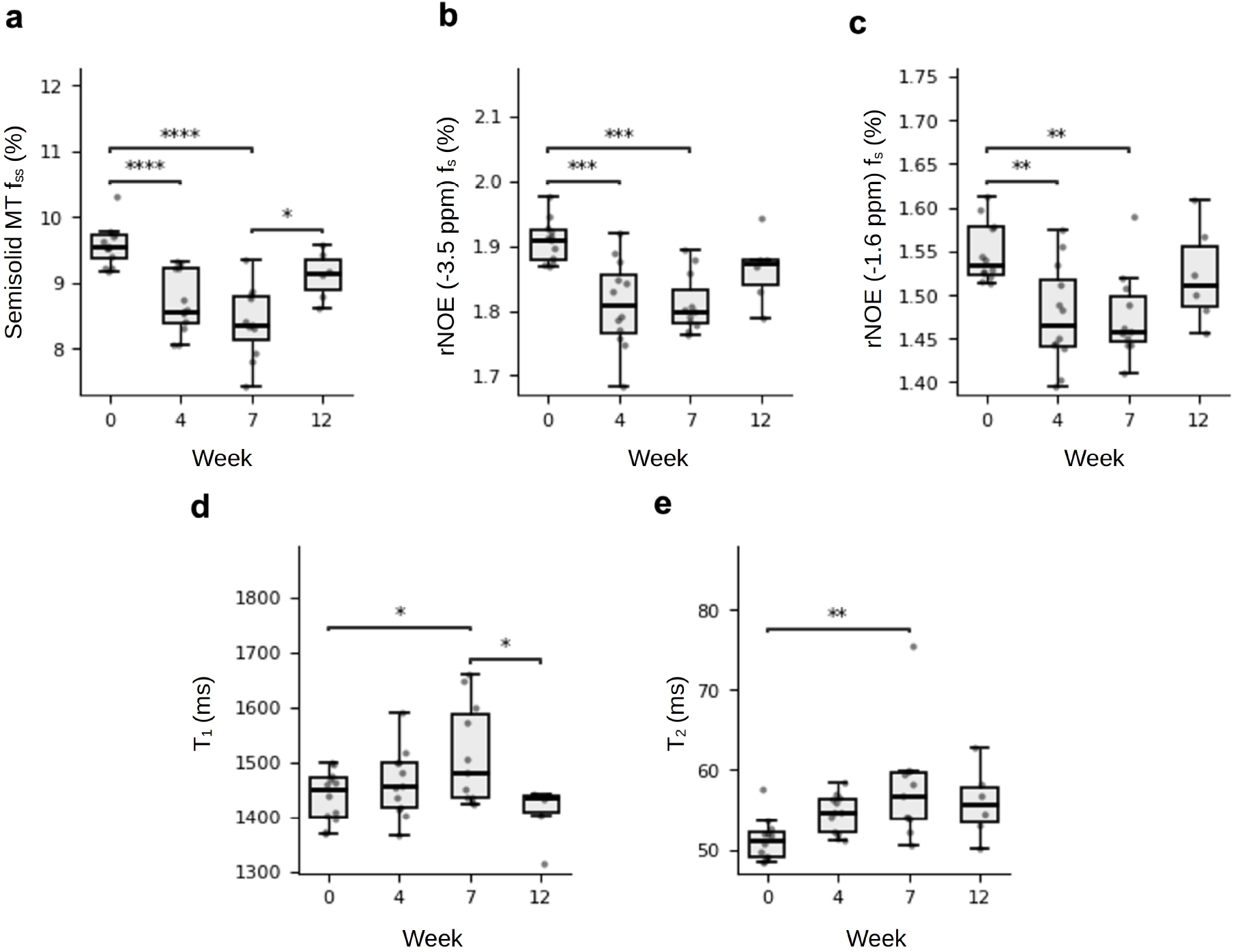
Statistical analysis of quantitative MRI parameters within the corpus callosum ROI across all four experimental time points using one-way analysis of variance (ANOVA) followed by Tukey’s HSD test. The examined parameters included the semisolid MT proton volume fraction (**a**), rNOE proton volume fraction at −3.5 ppm (**b**) and rNOE proton volume fraction at −1.6 ppm (**c**), water T_1_ (**d**), and T_2_ (**e**). *p<0.05; **p<0.01; ***p<0.001; ****p<0.0001.

As shown in **Figure. 5a**, there was a significant and progressive reduction in the semisolid MT proton volume fraction in the corpus callosum between baseline and week 4 (p < 0.0001) and between baseline and week 7 (p < 0.0001). However, the semisolid MT values increased significantly between weeks 7 and 12, after stopping the cuprizone diet, (p = 0.01), which is consistent with remyelination. The rNOE proton volume fractions (**Figure 5b,c**) were also significantly lower than baseline at both week 4 (p = 0.0002 and p = 0.005 for the rNOE at −3.5 ppm and −1.6 ppm, respectively) and week 7 (p = 0.0003 and p = 0.005 for the rNOE at −3.5 ppm and −1.6 ppm, respectively). Following withdrawal of cuprizone, both rNOE pools exhibited a general increase in proton volume fraction but the changes did not reach statistical significance.

There were significant increases in water T_1_ and T_2_ values in the corpus callosum between baseline and week 7 (p < 0.05), corresponding to the peak demyelination stage. Interestingly, there were no significant differences at week 4. Cessation of cuprizone treatment caused a statistically significant decrease in water T_1_ (p=0.02, **Figure 5d**, week 12 compared to week 7) but there was no corresponding significant difference in T_2_ values. Taken together, the analysis demonstrates the superior sensitivity of ST-MRF derived proton volume fractions to early-stage myelin damage, as they recognized the demyelination process at an earlier time-point than detected by conventional MRI contrast parameters.

Although quantitative T_1_ and T_2_ mapping is often used in MS research, the sensitivity to demyelination in the cuprizone model has been inconsistent across studies. Friesen et al. who conducted a cuprizone mouse model study at 7T^44^ reported significantly elevated T_1_ values in the corpus callosum at weeks 5 and 6 of cuprizone treatment, while T_2_ exhibited only a non-significant increase. Notably, that was an ex vivo study using separate cohorts of animals at individual endpoints rather than longitudinal in vivo imaging. In contrast, Hertanu et al. found no significant T_1_ changes during the demyelination phase in a 7T MRI cuprizone mouse study^45^ while Zinnhardt et al. reported a significant increase in T_2_ relaxation times at both 3 and 5 weeks of cuprizone treatment.^46^ In the present study, both T_1_ and T_2_ maps showed a significant change between baseline and week 7 - the peak demyelination timepoint. The inconsistencies across studies may be explained by two factors. First, the cited studies differ methodologically - including cuprizone dosage, sample size, experimental design, and imaging time points, thereby limiting a direct comparison. Second, more fundamentally, T_1_ and T_2_ relaxation times are influenced by multiple biological factors beyond myelin content alone. A recent meta-analysis of comparing quantitative MRI with histopathology in MS demonstrated moderate to strong correlations between T_1_ and T_2_ relaxation times and myelin density, while also identifying contributions from axonal injury and gliosis.^47^ These findings emphasize the need to employ multiple complementary specific biomarkers capable of disentangling distinct pathological processes associated with demyelination.

Our quantitative semisolid MT MRF findings are in accordance with previous MT-weighted reports that cuprizone feeding produces a noticeable decrease in MTR signal after demyelination, which then increases after remyelination. Notably, effects, obtained by using far saturation pulse frequency offsets, where CEST and rNOE contributions are expected to be negligible, primarily reflect changes in the semisolid macromolecular pool associated with myelin.^10,48-50^

Our findings are also in agreement with previously reported quantitative MT (qMT) studies. For example, Turati et al. reported a similar pattern for the qMT-derived proton pool volume fraction in a longitudinal 7T study using two mouse strains (male C57BL/6 and female SJL/J) fed cuprizone for 5 and 7 weeks respectively.^51^ The proton volume fraction in both strains declined over the course of the cuprizone diet relative to untreated controls. However, after cuprizone withdrawal the proton volume fraction of the C57BL/6 mice recovered by week 10, consistent with remyelination, while the SJL/J mice did not recover. The authors attributed this discrepancy to a combination of inherent strain differences in remyelination capacity and a shorter recovery window in SJL/J mice. The main limitation of this study was the >40 minute acquisition time for qMT (not including the T_1_ mapping time). A rapid qMT approach in human studies requires as little as 4 minutes for acquisition,^52^ but can quantify only a single proton pool (semisolid MT). In a comparable time of 5 minutes 45 seconds, the proposed ST-MRF framework jointly resolves f_ss_/f_s_ for three distinct molecular pools: semisolid MT, rNOE(−3.5 ppm), and rNOE(−1.6 ppm) simultaneously. Moreover, if semisolid MT quantification is deemed sufficient, it can be acquired within 105 seconds using the proposed MRF approach.

With respect to rNOE imaging at −3.5 ppm, Chen et al. showed a significant decrease in rNOE (−3.5 ppm) signal in the corpus callosum in a 3T longitudinal cuprizone mouse study.^16^ rNOE decreased significantly by week 8 of cuprizone diet compared to healthy mice, consistent with demyelination, and exhibited full recovery by week 14, consistent with remyelination. This is in agreement with our findings. The added value of MRF in this context is the ability to differentiate the effect of rNOE at −3.5 ppm from that of the MT, T_1_, and rNOE at −1.6 ppm.

Although the rNOE at −1.6 ppm has not been directly investigated in the context of MS or myelin, Wu et al. attributed this resonance to the choline methyl groups of phosphatidylcholines and showed that the amplitude correlates positively with cholesterol content in a glioma tumor model.^20^ Since myelin is among the most cholesterol-rich membranes in the body,^53^ these findings further support the potential of the rNOE (−1.6 ppm) as a biomarker of myelin integrity.

Notably, while the semisolid MT quantification framework allows the exchange rate to vary as a free parameter (**Figure S2** and **Table S1**), the rNOE analysis assumes a fixed proton exchange rate (of 16 s^-1^),^54^ consistent with accumulating evidence indicating that this exchange process exhibits minimal sensitivity to pH.^55-57^ Applying the pH weighted AACID approach to the same mice included in our study did not yield any significant effects across all timepoints (**Figure S3**), supporting the use of a fixed exchange rate in the rNOE modeling framework.

Amide proton transfer (APT)-weighted imaging, as analyzed via the MTR_asym_ value at +3.5 ppm relative to water, has recently been proposed as a potentially useful biomarker for monitoring myelination in a similar cuprizone mouse model.^58^ For comparison, we applied this metric on Z-spectra acquired on the same mice analyzed in our study (**Figure S4a**). While similar trends were obtained (MTR_asym_ increase on demyelination and decrease remyelination **Figure. S4c**), the effect did not reach statistical significance (p = 0.218, one-way ANOVA). While APT quantification was not included in the current MRF framework, the methodology could readily be expanded to incorporate APT related parameters, as previously described in other biomedical imaging applications.^27^ Interestingly, while clear rNOE effects are noticeable in the Z-spectra at both −3.5 ppm and −1.6 ppm (**Figure S4a**), longitudinal changes in the conventional MTR_asym_ metric at −1.6 ppm were considerably less pronounced (**Figure S4b**). This discrepancy may arise from confounding T_1_ dependent effects and potential contamination from opposing OH-CEST signals near +1.6 ppm.

### Histological Analysis

Three stains were used for histological analysis:

1. Myelin Basic Protein (MBP), an abundant myelin protein used to evaluate myelin integrity.^59^
2. Glial Fibrillary Acidic Protein (GFAP), a marker of astrocytes^60^ that become activated during the demyelination process in the cuprizone mouse model.^61^ This makes GFAP a useful indicator of demyelination.
3. DAPI (4′,6-diamidino-2-phenylindole), which stains cell nuclei regardless of cell type and was used as a counterstain to confirm tissue morphology and region-of-interest boundaries. Representative whole-slice images are shown in **Figure S5**, with a closer look at the corpus callosum region provided in **Figure 6**. Histological assessment was performed at baseline, week 7 (peak demyelination), and week 12 (recovery). The histological findings were in good agreement with the ST-MRF derived metrics. Specifically, there was a visible reduction in MBP intensity at week 7 relative to baseline, consistent with the significant decreases observed in semisolid MT and both rNOE pools over the same interval. This reduction was followed by a visual recovery in MBP signal at week 12, reflecting the recovery trend observed in the ST-MRF derived fractions (**Figure 4, Figure 5, Figure 7**). Furthermore, there was a marked increase in GFAP staining at week 7, reflecting the neuroinflammatory response known to accompany cuprizone-induced demyelination.^61,62^ Together, these histological findings provide independent validation that the ST-MRF derived parameters capture both myelin degeneration and subsequent recovery.

**Figure 6.**
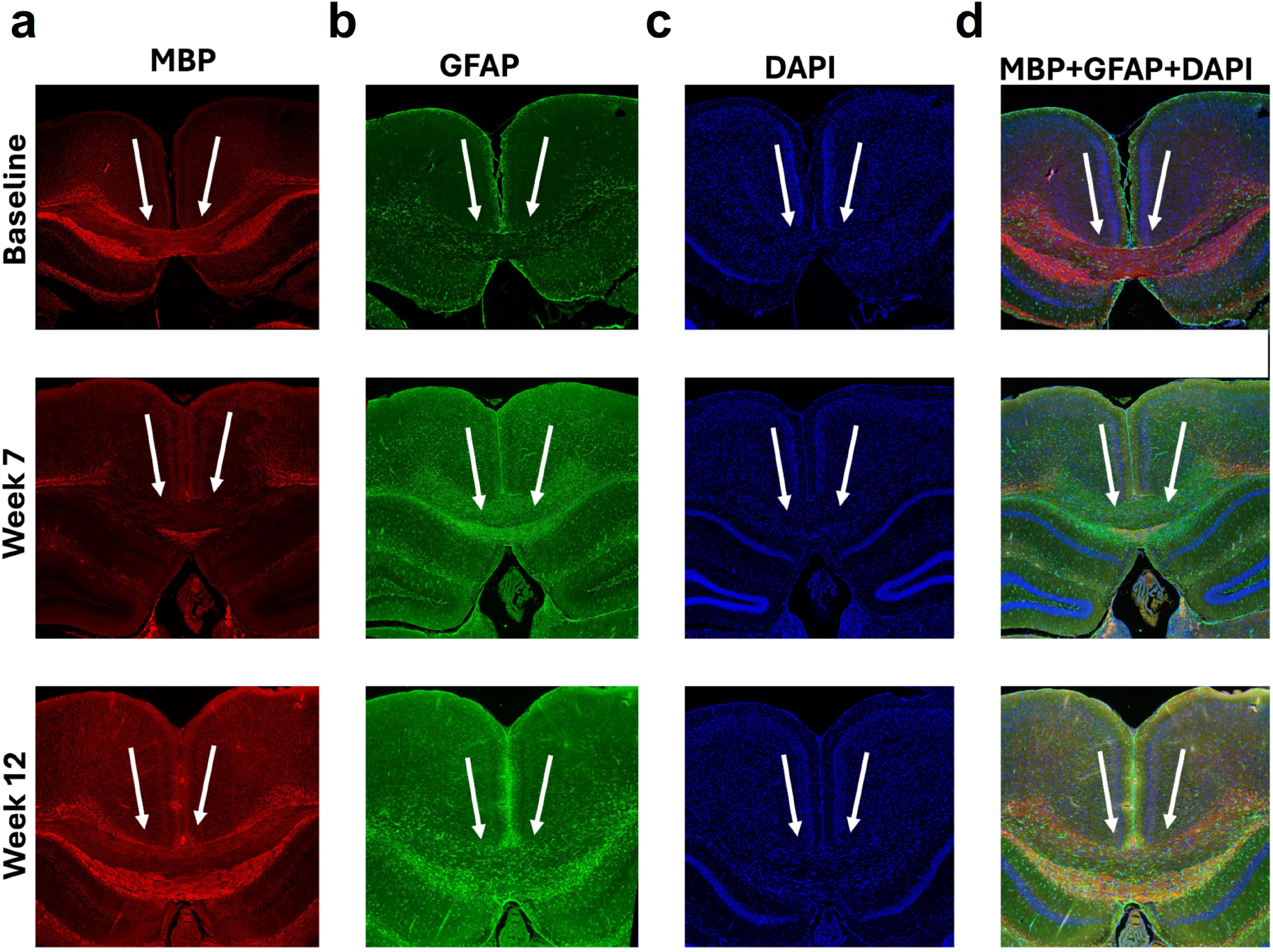
Histological images in three representative mice across three time points: baseline (**top**), week 7 of cuprizone feeding (**center**), and week 12 (five weeks after ceasing cuprizone feeding, **bottom**). Samples were stained for MBP (**a**), GFAP (**b**), and DAPI (**c**), with combined visualization shown in (**d**). Note the clear decrease in MBP signal at demyelination (week 7), followed by its increase at remyelination (week 12). White arrows point to the corpus callosum.

**Figure 7.**
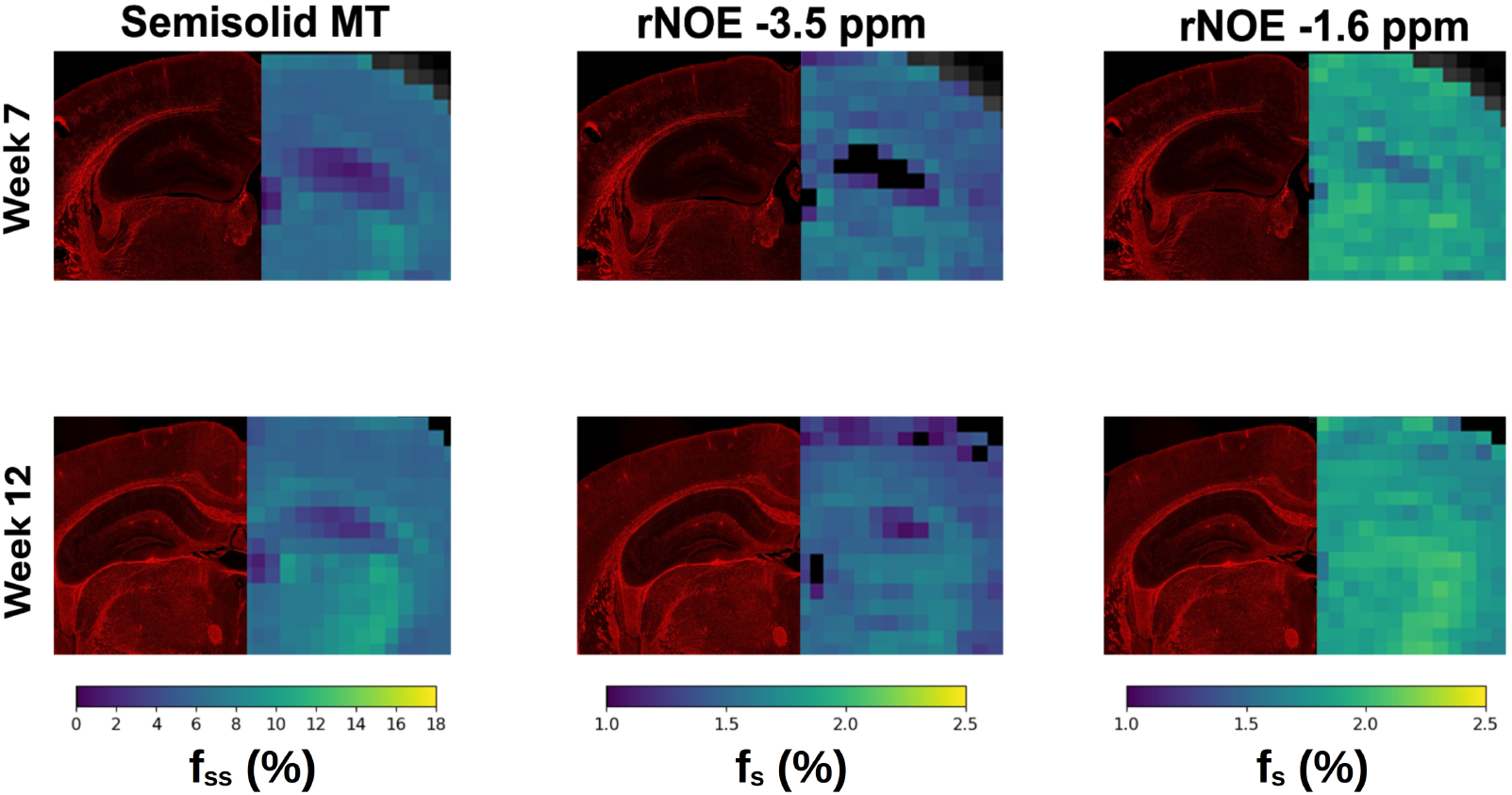
Side-by-side visualization of ST-MRF based proton volume fraction maps (right part of each fused image) alongside MBP histological images (left part of each fused image). Images are shown at peak demyelination (week 7 of cuprizone feeding, **top**) and remyelination (week 12, five weeks after ceasing cuprizone feeding, **bottom**) from three representative mice.

### Limitations and Future Work

This study has several limitations. First, the cuprizone model primarily recapitulates demyelination, one important aspect of MS pathology, but does not fully capture the complexity of the disease, which also involves processes such as neuroinflammation, gliosis, axonal injury, and immune cell infiltration.^63^ Consequently, the findings reported here should be interpreted within the specific context of toxin-induced demyelination. Second, this study was conducted with a 7T scanner. While this field strength is FDA approved for human imaging, such scanners are substantially less common than 3T scanners in clinical practice. However, the semisolid MT and rNOE proton pools investigated in this work lie in the slow exchange regime, and there is considerable evidence they are detectable in the human brain at 3T,^15,26^ suggesting a potential avenue for translation.

From an acquisition standpoint, the requirement for separately acquired T_1_ and T_2_ maps increases total examination time. While our experiments used standard relaxometry sequences, future human studies could leverage the very short water T_1_ and T_2_ MRF,^24^ which is already commercially available and FDA approved for selected scanner platforms. Such approaches could further reduce scan time while maintaining quantification accuracy.

Finally, the corpus callosum was selected as the primary region of interest because of previous reports that it is among the most severely affected white matter structures in the cuprizone model;^16,30,32,44,61,64^ however, our imaging protocol employed a single-slice acquisition centered on the body/isthmus corpus of the corpus callosum. However, we do appreciate that previous studies have reported regional differences in demyelination and recovery patterns within the corpus callosum, including the genu,^10^ as well as pathological changes in deeper gray matter structures located in more rostral brain regions.^49^ Therefore, future studies employing volumetric or multi-slice acquisitions will be important in order to evaluate the spatial heterogeneity of these effects and to assess the generalizability of the proposed biomarkers across the brain.

## Conclusions

This study introduces a rapid, quantitative, deep-learning-based ST-MRF framework capable of simultaneously isolating and quantifying multiple proton exchange parameters associated with myelin integrity in vivo. All three ST-MRF-derived parameters identified significant alterations in the corpus callosum at week 4, which is earlier than achieved by conventional T_1_- and T_2_- weighted imaging. Together, these results offer quantitative ST-MRF as a promising approach for the noninvasive assessment of myelin pathology and suggest potential utility as an early imaging biomarker for monitoring disease progression and therapeutic response in demyelinating disorders.

## Data Availability Statement

Phantom and mouse data will become available upon acceptance at https://github.com/momentum-laboratory/st-mrf-ms and Zenodo.

## Code Availability Statement

The code used in this work will become available upon acceptance at https://github.com/momentum-laboratory/st-mrf-ms and Zenodo. All ST-MRF acquisition protocols can be reproduced using the open MRI pulse sequences provided in https://osf.io/52bsg.^34^

## Author Contributions

R.B.C and O.P. designed the computational framework. M.R., R.B.C., and O.P. conceived the phantom and animal studies. R.B.C. and M.R. acquired the MRI data. R.B.C. performed the optimization, training, inference, and data analysis. R.B.C. wrote the manuscript. All authors reviewed and revised the manuscript. O.P. supervised the project.

## Acknowledgment

This work was supported by the Israel Science Foundation (grant No. 1030/25). This project was funded by the European Union (ERC, BabyMagnet, project no. 101115639). Views and opinions expressed are, however, those of the authors only and do not necessarily reflect those of the European Union or the European Research Council. Neither the European Union nor the granting authority can be held responsible for them.

## References

(1) Woo, M. S.; Engler, J. B.; Friese, M. A. The Neuropathobiology of Multiple Sclerosis. Nat. Rev. Neurosci. 2024, 25 (7), 493–513. 10.1038/s41583-024-00823-z.

(2) Selmaj, K.; Cree, B. A. C.; Barnett, M.; Thompson, A.; Hartung, H.-P. Multiple Sclerosis: Time for Early Treatment with High-Efficacy Drugs. J. Neurol. 2024, 271 (1), 105–115. 10.1007/s00415-023-11969-8.

(3) Blaschke, S. J.; Ellenberger, D.; Flachenecker, P.; Hellwig, K.; Paul, F.; Pöhlau, D.; Kleinschnitz, C.; Rommer, P. S.; Rueger, M. A.; Zettl, U. K.; Stahmann, A.; Warnke, C. Time to Diagnosis in Multiple Sclerosis: Epidemiological Data from the German Multiple Sclerosis Registry. Mult. Scler. J. 2022, 28 (6), 865–871. 10.1177/13524585211039753.

(4) Montalban, X.; Lebrun-Frénay, C.; Oh, J.; Arrambide, G.; Moccia, M.; Pia Amato, M.; Amezcua, L.; Banwell, B.; Bar-Or, A.; Barkhof, F.; Butzkueven, H.; Ciccarelli, O.; Chataway, J.; Cohen, J. A.; Comi, G.; Correale, J.; Deisenhammer, F.; Filippi, M.; Fiol, J.; Freedman,M. S.; Fujihara, K.; Granziera, C.; Green, A. J.; Hartung, H.-P.; Hellwig, K.; Kappos, L.; Kimbrough, D.; Killestein, J.; Lublin, F.; Marignier, R.; Ann Marrie, R.; Miller, A.; Otero-Romero, S.; Ontaneda, D.; Ramanathan, S.; Reich, D.; Rocca, M. A.; Rovira, À.; Saidha, S.; Salter, A.; Sastre-Garriga, J.; Saylor, D.; Solomon, A. J.; Sormani, M. P.; Stankoff, B.; Tintore, M.; Tremlett, H.; Van Der Walt, A.; Viswanathan, S.; Wiendl, H.; Wildemann, B.; Yamout, B.; Zaratin, P.; Calabresi, P. A.; Coetzee, T.; Thompson, A. J. Diagnosis of Multiple Sclerosis: 2024 Revisions of the McDonald Criteria. Lancet Neurol. 2025, 24 (10), 850–865. 10.1016/S1474-4422(25)00270-4.

(5) Solomon, A. J.; Arrambide, G.; Brownlee, W. J.; Flanagan, E. P.; Amato, M. P.; Amezcua, L.; Banwell, B. L.; Barkhof, F.; Corboy, J. R.; Correale, J.; Fujihara, K.; Graves, J.; Harnegie,M. P.; Hemmer, B.; Lechner-Scott, J.; Marrie, R. A.; Newsome, S. D.; Rocca, M. A.; Royal, W.; Waubant, E. L.; Yamout, B.; Cohen, J. A. Differential Diagnosis of Suspected Multiple Sclerosis: An Updated Consensus Approach. Lancet Neurol. 2023, 22 (8), 750–768. 10.1016/S1474-4422(23)00148-5.

(6) Jones, K. M.; Pollard, A. C.; Pagel, M. D. Clinical Applications of Chemical Exchange Saturation Transfer (CEST) MRI. J. Magn. Reson. Imaging 2018, 47 (1), 11–27. 10.1002/jmri.25838.

(7) Van Zijl, P. C. M.; Yadav, N. N. Chemical Exchange Saturation Transfer (CEST): What Is in a Name and What Isn’t? Magn. Reson. Med. 2011, 65 (4), 927–948. 10.1002/mrm.22761.

(8) Dortch, R. D.; Moore, J.; Li, K.; Jankiewicz, M.; Gochberg, D. F.; Hirtle, J. A.; Gore, J. C.; Smith, S. A. Quantitative Magnetization Transfer Imaging of Human Brain at 7 T. NeuroImage 2013, 64, 640–649. 10.1016/j.neuroimage.2012.08.047.

(9) Van Gelderen, P.; Duyn, J. H. White Matter Intercompartmental Water Exchange Rates Determined from Detailed Modeling of the Myelin Sheath. Magn. Reson. Med. 2019, 81 (1), 628–638. 10.1002/mrm.27398.

(10) Tagge, I.; O’Connor, A.; Chaudhary, P.; Pollaro, J.; Berlow, Y.; Chalupsky, M.; Bourdette, D.; Woltjer, R.; Johnson, M.; Rooney, W. Spatio-Temporal Patterns of Demyelination and Remyelination in the Cuprizone Mouse Model. PLOS ONE 2016, 11 (4), e0152480. 10.1371/journal.pone.0152480.

(11) York, E. N.; Thrippleton, M. J.; Meijboom, R.; Hunt, D. P. J.; Waldman, A. D. Quantitative Magnetization Transfer Imaging in Relapsing-Remitting Multiple Sclerosis: A Systematic Review and Meta-Analysis. Brain Commun. 2022, 4 (2), fcac088. 10.1093/braincomms/fcac088.

(12) Guglielmetti, C.; Boucneau, T.; Cao, P.; Van Der Linden, A.; Larson, P. E. Z.; Chaumeil,M. M. Longitudinal Evaluation of Demyelinated Lesions in a Multiple Sclerosis Model UsingUltrashort Echo Time Magnetization Transfer (UTE-MT) Imaging. NeuroImage 2020, 208, 116415. 10.1016/j.neuroimage.2019.116415.

(13) Zhang, H.; Wang, Z.; Zeng, S.; Wang, J.; Cai, P.; Mak, H. K. F.; Chan, K. H.; Qiu, B.; Huang, J. Quasi-Steady-State CEST for Rapid and Quantitative Lesion Detection in Multiple Sclerosis at 3T. Brain Res. Bull. 2026, 237, 111774. 10.1016/j.brainresbull.2026.111774.

(14) Zhou, Y.; Bie, C.; Van Zijl, P. C. M.; Yadav, N. N. The Relayed Nuclear Overhauser Effect in Magnetization Transfer and Chemical Exchange Saturation Transfer MRI. NMR Biomed. 2023, 36 (6), e4778. 10.1002/nbm.4778.

(15) Huang, J.; Xu, J.; Lai, J. H. C.; Chen, Z.; Lee, C. Y.; Mak, H. K. F.; Chan, K. H.; Chan, K.W. Y. Relayed Nuclear Overhauser Effect Weighted (rNOEw) Imaging Identifies Multiple Sclerosis. NeuroImage Clin. 2021, 32, 102867. 10.1016/j.nicl.2021.102867.

(16) Chen, Z.; Huang, J.; Lai, J. H. C.; Tse, K.; Xu, J.; Chan, K. W. Y. Chemical Exchange Saturation Transfer MRI Detects Myelin Changes in Cuprizone Mouse Model at 3T. NMR Biomed. 2023, 36 (9), e4937. 10.1002/nbm.4937.

(17) Zhang, X.-Y.; Wang, F.; Afzal, A.; Xu, J.; Gore, J. C.; Gochberg, D. F.; Zu, Z. A New NOE-Mediated MT Signal at around −1.6ppm for Detecting Ischemic Stroke in Rat Brain. Magn. Reson. Imaging 2016, 34 (8), 1100–1106. 10.1016/j.mri.2016.05.002.

(18) Zhang, X.-Y.; Wang, F.; Jin, T.; Xu, J.; Xie, J.; Gochberg, D. F.; Gore, J. C.; Zu, Z. MR Imaging of a Novel NOE-Mediated Magnetization Transfer with Water in Rat Brain at 9.4 T. Magn. Reson. Med. 2017, 78 (2), 588–597. 10.1002/mrm.26396.

(19) Viswanathan, M.; Kurmi, Y.; Zu, Z. Nuclear Overhauser Enhancement Imaging at −1.6 Ppm in Rat Brain at 4.7T. Magn. Reson. Med. 2024, 91 (2), 615–629. 10.1002/mrm.29896.

(20) Wu, Q.-X.; Liu, H.-Q.; Wang, Y.-J.; Chen, T.-C.; Wei, Z.-Y.; Chang, J.-H.; Chen, T.-H.; Seema, J.; Lin, E. C. Chemical Exchange Saturation Transfer (CEST) Signal at −1.6 Ppm and Its Application for Imaging a C6 Glioma Model. Biomedicines 2022, 10 (6), 1220. 10.3390/biomedicines10061220.

(21) Morini, M. A.; Pedroni, V. I. Role of Lipid Composition on the Mechanical and Biochemical Vulnerability of Myelin and Its Implications for Demyelinating Disorders. Biophysica 2025, 5 (4), 44. 10.3390/biophysica5040044.

(22) Liu, G.; Song, X.; Chan, K. W. Y.; McMahon, M. T. Nuts and Bolts of Chemical Exchange Saturation Transfer MRI. NMR Biomed. 2013, 26 (7), 810–828. 10.1002/nbm.2899.

(23) Cohen, O.; Yu, V. Y.; Tringale, K. R.; Young, R. J.; Perlman, O.; Farrar, C. T.; Otazo, R. CEST MR Fingerprinting (CEST-MRF) for Brain Tumor Quantification Using EPI Readout and Deep Learning Reconstruction. Magn. Reson. Med. 2023, 89 (1), 233–249. 10.1002/mrm.29448.

(24) Ma, D.; Gulani, V.; Seiberlich, N.; Liu, K.; Sunshine, J. L.; Duerk, J. L.; Griswold, M. A. Magnetic Resonance Fingerprinting. Nature 2013, 495 (7440), 187–192. 10.1038/nature11971.

(25) Singh, M.; Kang, B.; Mahmud, S. Z.; Van Zijl, P.; Zhou, J.; Heo, H. Saturation Transfer MR Fingerprinting for Magnetization Transfer Contrast and Chemical Exchange Saturation Transfer Quantification. Magn. Reson. Med. 2025, 94 (3), 993–1009. 10.1002/mrm.30532.

(26) Power, I.; Rivlin, M.; Shmuely, H.; Zaiss, M.; Navon, G.; Perlman, O. In Vivo Mapping of the Chemical Exchange Relayed Nuclear Overhauser Effect Using Deep Magnetic Resonance Fingerprinting. iScience 2024, 27 (11), 111209. 10.1016/j.isci.2024.111209.

(27) Perlman, O.; Ito, H.; Herz, K.; Shono, N.; Nakashima, H.; Zaiss, M.; Chiocca, E. A.; Cohen, O.; Rosen, M. S.; Farrar, C. T. Quantitative Imaging of Apoptosis FollowingOncolytic Virotherapy by Magnetic Resonance Fingerprinting Aided by Deep Learning. Nat. Biomed. Eng. 2021, 6 (5), 648–657. 10.1038/s41551-021-00809-7.

(28) Shmuely, H.; Rivlin, M.; Perlman, O. Quantitative Multi-Metabolite Imaging of Parkinson’s Disease Using AI Boosted Molecular MRI. Npj Imaging 2025, 3 (1), 66. 10.1038/s44303-025-00130-x.

(29) Prasuhn, J.; Singh, M.; Mahmud, S. Z.; Yadav, N. N.; Dawson, T. M.; Mills, K. A.; Zijl, P. V.; Heo, H.-Y. Deep-Learning Saturation Transfer Magnetic Resonance Fingerprinting (ST-MRF) in Patients with Parkinson’s Disease. NeuroImage 2026, 334, 121975. 10.1016/j.neuroimage.2026.121975.

(30) Matsushima, G. K.; Morell, P. The Neurotoxicant, Cuprizone, as a Model to Study Demyelination and Remyelination in the Central Nervous System. Brain Pathol. 2001, 11 (1), 107–116. 10.1111/j.1750-3639.2001.tb00385.x.

(31) Ma, Y.-J.; Jang, H.; Chang, E. Y.; Hiniker, A.; Head, B. P.; Lee, R. R.; Corey-Bloom, J.; Bydder, G. M.; Du, J. Ultrashort Echo Time (UTE) Magnetic Resonance Imaging of Myelin: Technical Developments and Challenges. Quant. Imaging Med. Surg. 2020, 10 (6), 1186–1203. 10.21037/qims-20-541.

(32) Hiremath, M. M.; Saito, Y.; Knapp, G. W.; Ting, J. P.-Y.; Suzuki, K.; Matsushima, G. K. Microglial/Macrophage Accumulation during Cuprizone-Induced Demyelination in C57BL/6 Mice. J. Neuroimmunol. 1998, 92 (1–2), 38–49. 10.1016/S0165-5728(98)00168-4.

(33) Vladimirov, N.; Cohen, O.; Heo, H.-Y.; Zaiss, M.; Farrar, C. T.; Perlman, O. Quantitative Molecular Imaging Using Deep Magnetic Resonance Fingerprinting. Nat. Protoc. 2025, 20 (10), 3024–3054. 10.1038/s41596-025-01152-w.

(34) Cohen, O.; Huang, S.; McMahon, M. T.; Rosen, M. S.; Farrar, C. T. Rapid and Quantitative Chemical Exchange Saturation Transfer (CEST) Imaging with Magnetic Resonance Fingerprinting (MRF). Magn. Reson. Med. 2018, 80 (6), 2449–2463. 10.1002/mrm.27221.

(35) McVicar, N.; Li, A. X.; Gonçalves, D. F.; Bellyou, M.; Meakin, S. O.; Prado, M. A.; Bartha,R. Quantitative Tissue Ph Measurement during Cerebral Ischemia Using Amine and Amide Concentration-Independent Detection (AACID) with MRI. J. Cereb. Blood Flow Metab. 2014, 34 (4), 690–698. 10.1038/jcbfm.2014.12.

(36) Lein, E. S.; Hawrylycz, M. J.; Ao, N.; Ayres, M.; Bensinger, A.; Bernard, A.; Boe, A. F.; Boguski, M. S.; Brockway, K. S.; Byrnes, E. J.; Chen, L.; Chen, L.; Chen, T.-M.; Chi Chin, M.; Chong, J.; Crook, B. E.; Czaplinska, A.; Dang, C. N.; Datta, S.; Dee, N. R.; Desaki, A. L.; Desta, T.; Diep, E.; Dolbeare, T. A.; Donelan, M. J.; Dong, H.-W.; Dougherty, J. G.; Duncan,B. J.; Ebbert, A. J.; Eichele, G.; Estin, L. K.; Faber, C.; Facer, B. A.; Fields, R.; Fischer, S. R.; Fliss, T. P.; Frensley, C.; Gates, S. N.; Glattfelder, K. J.; Halverson, K. R.; Hart, M. R.; Hohmann, J. G.; Howell, M. P.; Jeung, D. P.; Johnson, R. A.; Karr, P. T.; Kawal, R.; Kidney,J. M.; Knapik, R. H.; Kuan, C. L.; Lake, J. H.; Laramee, A. R.; Larsen, K. D.; Lau, C.; Lemon, T. A.; Liang, A. J.; Liu, Y.; Luong, L. T.; Michaels, J.; Morgan, J. J.; Morgan, R. J.; Mortrud, M. T.; Mosqueda, N. F.; Ng, L. L.; Ng, R.; Orta, G. J.; Overly, C. C.; Pak, T. H.; Parry, S. E.; Pathak, S. D.; Pearson, O. C.; Puchalski, R. B.; Riley, Z. L.; Rockett, H. R.; Rowland, S. A.; Royall, J. J.; Ruiz, M. J.; Sarno, N. R.; Schaffnit, K.; Shapovalova, N. V.; Sivisay, T.; Slaughterbeck, C. R.; Smith, S. C.; Smith, K. A.; Smith, B. I.; Sodt, A. J.; Stewart, N. N.; Stumpf, K.-R.; Sunkin, S. M.; Sutram, M.; Tam, A.; Teemer, C. D.; Thaller, C.; Thompson, C. L.; Varnam, L. R.; Visel, A.; Whitlock, R. M.; Wohnoutka, P. E.; Wolkey,C. K.; Wong, V. Y.; Wood, M.; Yaylaoglu, M. B.; Young, R. C.; Youngstrom, B. L.; Feng Yuan, X.; Zhang, B.; Zwingman, T. A.; Jones, A. R. Genome-Wide Atlas of Gene Expression in the Adult Mouse Brain. Nature 2007, 445 (7124), 168–176. 10.1038/nature05453.

(37) Seabold, S.; Perktold, J. Statsmodels: Econometric and Statistical Modeling with Python; Austin, Texas, 2010; pp 92–96. 10.25080/Majora-92bf1922-011.

(38) Morini, M. A.; Pedroni, V. I. Role of Lipid Composition on the Mechanical and Biochemical Vulnerability of Myelin and Its Implications for Demyelinating Disorders. Biophysica 2025, 5 (4), 44. 10.3390/biophysica5040044.

(39) Kim, H.-Y.; Huang, B. X.; Spector, A. A. Phosphatidylserine in the Brain: Metabolism and Function. Prog. Lipid Res. 2014, 56, 1–18. 10.1016/j.plipres.2014.06.002.

(40) Pousinis, P.; Begou, O.; Boziki, M. K.; Grigoriadis, N.; Theodoridis, G.; Gika, H. Recent Advances in Metabolomics and Lipidomics Studies in Human and Animal Models of Multiple Sclerosis. Metabolites 2024, 14 (10), 545. 10.3390/metabo14100545.

(41) Wheeler, D.; Bandaru, V. V. R.; Calabresi, P. A.; Nath, A.; Haughey, N. J. A Defect of Sphingolipid Metabolism Modifies the Properties of Normal Appearing White Matter in Multiple Sclerosis. Brain 2008, 131 (11), 3092–3102. 10.1093/brain/awn190.

(42) Boggs, J. M.; Rangaraj, G.; Dicko, A. Effect of Phosphorylation of Phosphatidylinositol on Myelin Basic Protein-Mediated Binding of Actin Filaments to Lipid Bilayers in Vitro. Biochim. Biophys. Acta BBA - Biomembr. 2012, 1818 (9), 2217–2227. 10.1016/j.bbamem.2012.04.006.

(43) Perlman, O.; Zhu, B.; Zaiss, M.; Rosen, M. S.; Farrar, C. T. An End-to-end AI-based Framework for Automated Discovery of Rapid CEST/MT MRI Acquisition Protocols and Molecular Parameter Quantification (AutoCEST). Magn. Reson. Med. 2022, 87 (6), 2792–2810. 10.1002/mrm.29173.

(44) Friesen, E.; Sheft, M.; Hari, K.; Palmer, V.; Zhu, S.; Herrera, S.; Buist, R.; Jiang, D.; Li, X.-M.; Del Bigio, Marc. R.; Thiessen, J. D.; Martin, M. Quantitative Analysis of Early White Matter Damage in Cuprizone Mouse Model of Demyelination Using 7.0 T MRI Multiparametric Approach. ASN Neuro 2024, 16 (1), 2404366. 10.1080/17590914.2024.2404366.

(45) Hertanu, A.; Soustelle, L.; Buron, J.; Le Priellec, J.; Cayre, M.; Le Troter, A.; Prevost, V. H.; Ranjeva, J.-P.; Varma, G.; Alsop, D. C.; Durbec, P.; Girard, O. M.; Duhamel, G. Inhomogeneous Magnetization Transfer (ihMT) Imaging in the Acute Cuprizone Mouse Model of Demyelination/Remyelination. NeuroImage 2023, 265, 119785. 10.1016/j.neuroimage.2022.119785.

(46) Zinnhardt, B.; Belloy, M.; Fricke, I. B.; Orije, J.; Guglielmetti, C.; Hermann, S.; Wagner, S.; Schäfers, M.; Van Der Linden, A.; Jacobs, A. H. Molecular Imaging of Immune Cell Dynamics During De- and Remyelination in the Cuprizone Model of Multiple Sclerosis by [18 F]DPA-714 PET and MRI. Theranostics 2019, 9 (6), 1523–1537. 10.7150/thno.32461.

(47) Shen, T.; Sheriff, S.; Qu, Y.; Gupta, V. K.; Graham, S. L.; Klistorner, A.; Jia, H.; Sun, X.; You, Y. Correlations between Postmortem Quantitative MRI Parameters and Demyelination, Axonal Loss and Gliosis in Multiple Sclerosis: A Systematic Review and Meta-Analysis. Brain Imaging Behav. 2025, 19 (2), 323–335. 10.1007/s11682-025-00971-5.

(48) Zaaraoui, W.; Deloire, M.; Merle, M.; Girard, C.; Raffard, G.; Biran, M.; Inglese, M.; Petry, K. G.; Gonen, O.; Brochet, B.; Franconi, J.-M.; Dousset, V. Monitoring Demyelination and Remyelination by Magnetization Transfer Imaging in the Mouse Brain at 9.4 T. Magn. Reson. Mater. Phys. Biol. Med. 2008, 21 (5), 357–362. 10.1007/s10334-008-0141-3.

(49) Fjær, S.; Bø, L.; Lundervold, A.; Myhr, K.-M.; Pavlin, T.; Torkildsen, Ø.; Wergeland, S. Deep Gray Matter Demyelination Detected by Magnetization Transfer Ratio in the Cuprizone Model. PLoS ONE 2013, 8 (12), e84162. 10.1371/journal.pone.0084162.

(50) Han, X.; Chen, J.; Liu, Z.; Liu, J.; Lin, M.; Wang, N. Detecting Early Brain Susceptibility Changes before Demyelination in Cuprizone Mouse Model Using Quantitative Susceptibility Mapping (QSM). NeuroImage 2026, 332, 121922. 10.1016/j.neuroimage.2026.121922.

(51) Turati, L.; Moscatelli, M.; Mastropietro, A.; Dowell, N. G.; Zucca, I.; Erbetta, A.; Cordiglieri, C.; Brenna, G.; Bianchi, B.; Mantegazza, R.; Cercignani, M.; Baggi, F.; Minati, L. In Vivo Quantitative Magnetization Transfer Imaging Correlates with Histology during De- and Remyelination in Cuprizone-treated Mice. NMR Biomed. 2015, 28 (3), 327–337. 10.1002/nbm.3253.

(52) Assländer, J.; Mao, A.; Marchetto, E.; Beck, E. S.; Rosa, F. L.; Charlson, R. W.; Shepherd, T. M.; Flassbeck, S. Unconstrained Quantitative Magnetization Transfer Imaging: Disentangling T1 of the Free and Semi-Solid Spin Pools. Imaging Neurosci. 2024, 2, imag– 2–00177. 10.1162/imag_a_00177.

(53) Kister, A.; Kister, I. Overview of Myelin, Major Myelin Lipids, and Myelin-Associated Proteins. Front. Chem. 2023, 10, 1041961. 10.3389/fchem.2022.1041961.

(54) Van Zijl, P. C. M.; Lam, W. W.; Xu, J.; Knutsson, L.; Stanisz, G. J. Magnetization Transfer Contrast and Chemical Exchange Saturation Transfer MRI. Features and Analysis of the Field-Dependent Saturation Spectrum. NeuroImage 2018, 168, 222–241. 10.1016/j.neuroimage.2017.04.045.

(55) Jin, T.; Wang, P.; Zong, X.; Kim, S. MR Imaging of the Amide-proton Transfer Effect and the pH-insensitive Nuclear Overhauser Effect at 9.4 T. Magn. Reson. Med. 2013, 69 (3), 760–770. 10.1002/mrm.24315.

(56) Jin, T.; Kim, S. Role of Chemical Exchange on the Relayed Nuclear Overhauser Enhancement Signal in Saturation Transfer MRI. Magn. Reson. Med. 2022, 87 (1), 365–376. 10.1002/mrm.28961.

(57) Zhao, Y.; Sun, C.; Zu, Z. Assignment of Molecular Origins of NOE Signal at −3.5 Ppm in the Brain. Magn. Reson. Med. 2023, 90 (2), 673–685. 10.1002/mrm.29643.

(58) Lee, D.-W.; Heo, H.; Woo, D.-C.; Kim, J. K.; Lee, D.-H. Amide Proton Transfer-Weighted 7-T MRI Contrast of Myelination after Cuprizone Administration. Radiology 2021, 299 (2), 428–434. 10.1148/radiol.2021203766.

(59) Boggs, J. M. Myelin Basic Protein: A Multifunctional Protein. Cell. Mol. Life Sci. 2006, 63(17), 1945–1961. 10.1007/s00018-006-6094-7.

(60) Yang, Z.; Wang, K. K. W. Glial Fibrillary Acidic Protein: From Intermediate Filament Assembly and Gliosis to Neurobiomarker. Trends Neurosci. 2015, 38 (6), 364–374. 10.1016/j.tins.2015.04.003.

(61) Kipp, M. Astrocytes: Lessons Learned from the Cuprizone Model. Int. J. Mol. Sci. 2023,24 (22), 16420. 10.3390/ijms242216420.

(62) Castillo-Rodriguez, M. D. L. A.; Gingele, S.; Schröder, L.-J.; Möllenkamp, T.; Stangel, M.; Skripuletz, T.; Gudi, V. Astroglial and Oligodendroglial Markers in the Cuprizone Animal Model for De- and Remyelination. Histochem. Cell Biol. 2022, 158 (1), 15–38. 10.1007/s00418-022-02096-y.

(63) Compston, A.; Coles, A. Multiple Sclerosis. The Lancet 2002, 359 (9313), 1221–1231. 10.1016/S0140-6736(02)08220-X.

(64) Thiessen, J. D.; Zhang, Y.; Zhang, H.; Wang, L.; Buist, R.; Del Bigio, M. R.; Kong, J.; Li, X.; Martin, M. Quantitative MRI and Ultrastructural Examination of the Cuprizone Mouse Model of Demyelination. NMR Biomed. 2013, 26 (11), 1562–1581. 10.1002/nbm.2992.

