## supporting information for "Quantitative Semisolid Magnetization Transfer and Relayed Nuclear Overhauser Effect Imaging in a Multiple Sclerosis Mouse Model Using Deep Magnetic Resonance Fingerprinting"

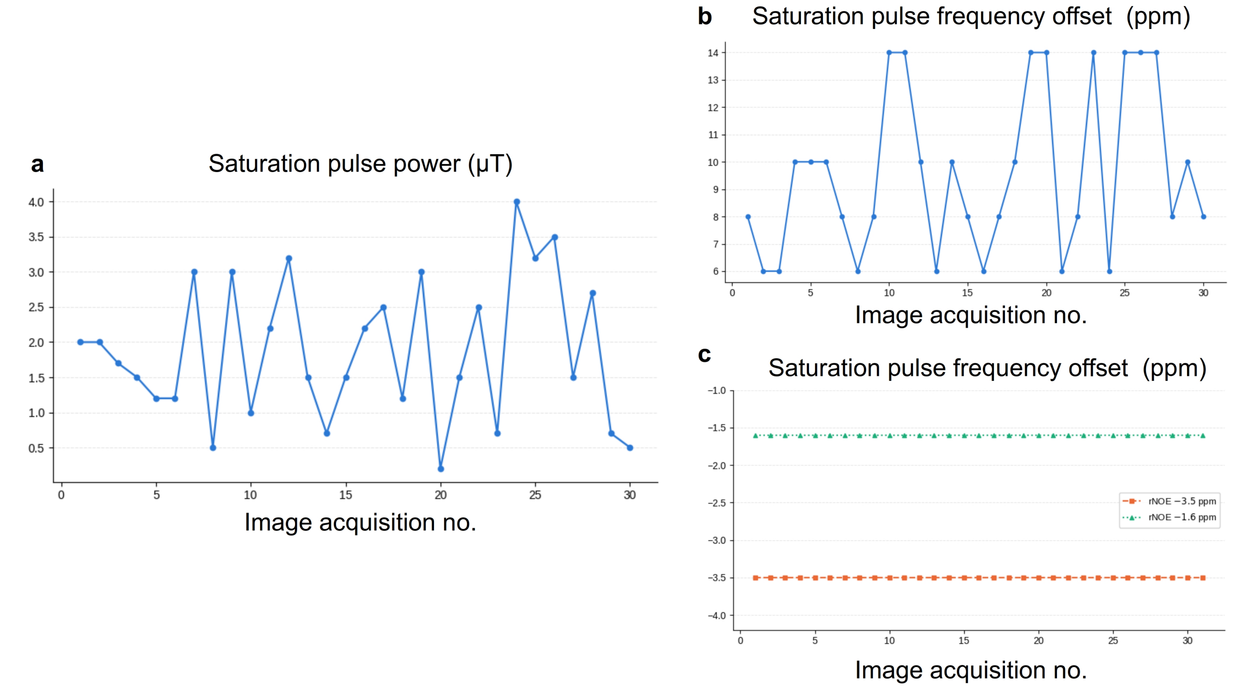


**Figure S1**. Acquisition parameters used in the MRF protocols. **a**. The same 0-4 µT saturation pulse power series[1] was used for encoding all proton pools, with an $M_0$ (no saturation, $B_1$=0) image added at the beginning of the rNOE protocols.[2] The saturation pulse frequency offset was varied between 6-14 ppm for encoding the semisolid MT pool[3] (**b**), or fixed at -3.5 ppm or -1.6 ppm for encoding the rNOE proton pools (**c**).

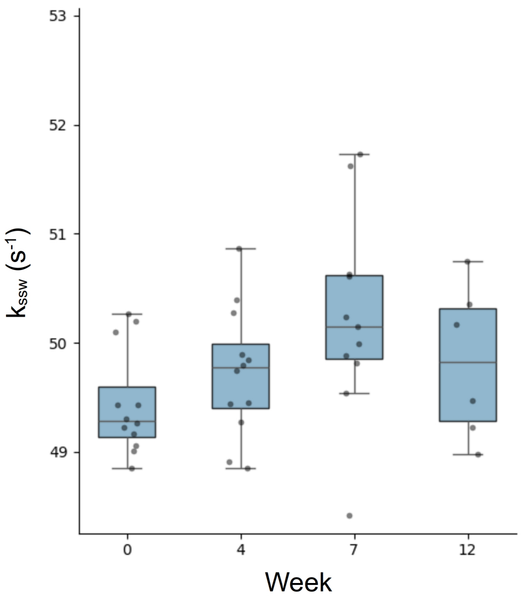


**Figure S2**. Longitudinal MT $k_{ssw}$ in the corpus callosum of the studied mice. One-way ANOVA showed no significant overall effect ($p = 0.06$).

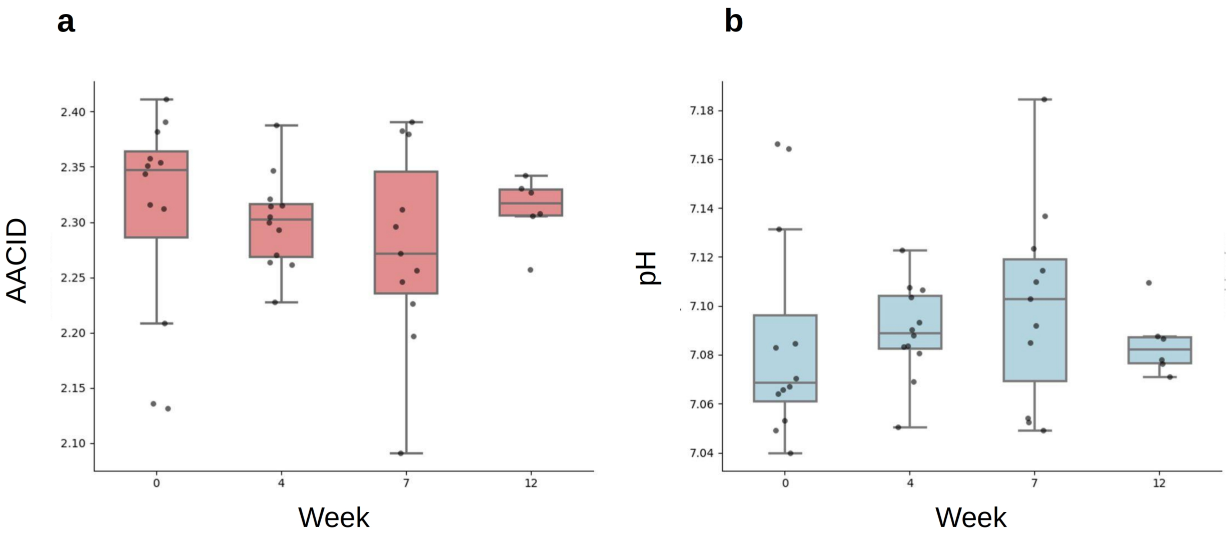


**Figure S3**. Longitudinal AACID (**a**) and pH (**b**) values calculated at the corpus callosum ROI. One way ANOVA in both cases did not reach statistical significance (p=0.736).

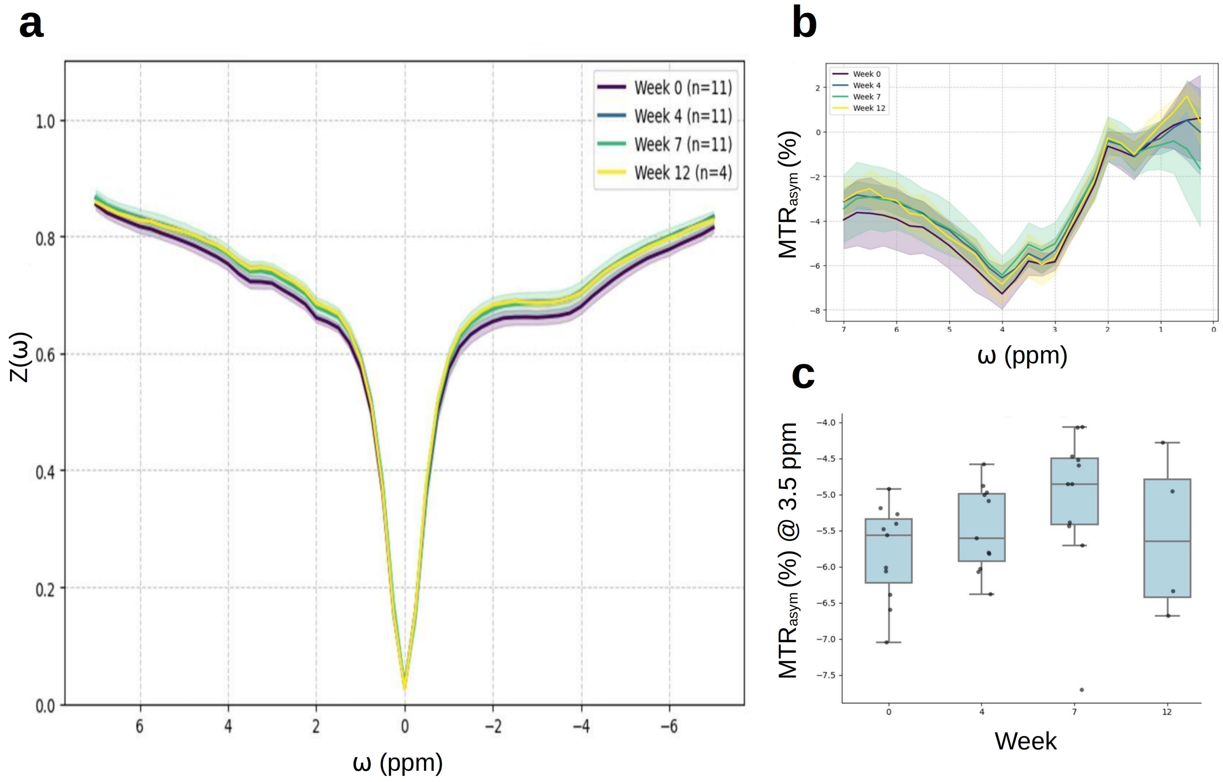


**Figure S4.** Longitudinal analysis of the Z-spectra (**a**), MTR asymmetry across all frequencies (**b**) and at $MTR_{asym}$ at 3.5 ppm (**c**), at the corpus callosum region of interest (ROI). Shaded regions in (**a-b**) represent the standard deviation.

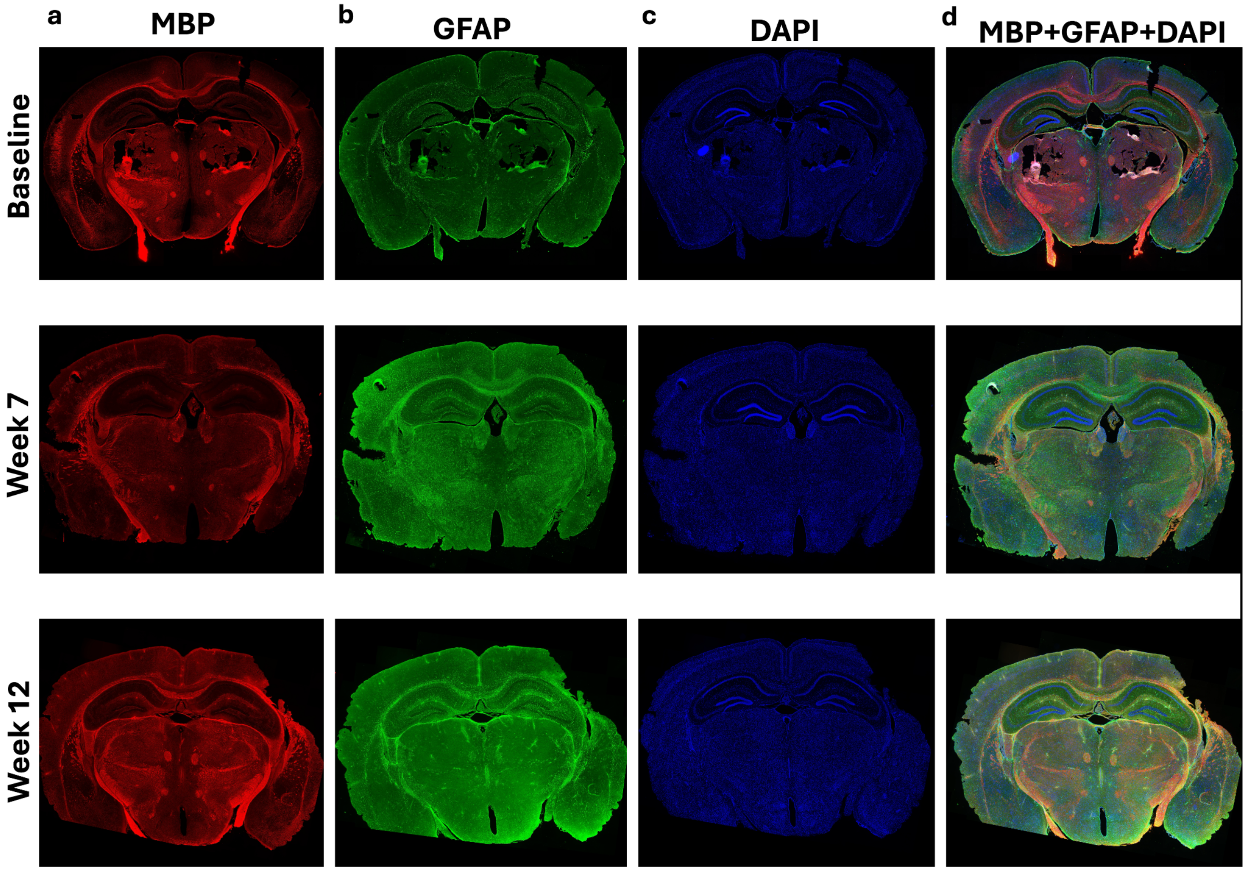


**Figure S5**. Histological images in three representative mice across three time points: baseline (top), week 7 of cuprizone feeding (center), and week 12 (five weeks after ceasing cuprizone feeding, bottom). Staining was performed using MBP (**a**), GFAP (**b**), and DAPI (**c**), with combined visualization shown in (**d**). Note the clear decrease in MBP signal at demyelination (week 7), followed by its increase at remyelination (week 12).

| MT dictionary (for training the first network) | | |
|---|---|---|
| Water | $T_1$ (ms) | 1200:100:2300* |
| | $T_2$ (ms) | 40:10:110 |
| MT | $T_1$ (ms) | Fixed to water |
| | $T_2$ (ms) | 0.04 |
| | Chemical shift (ppm) | -2.5 |
| | $f_{ss}$ (%) | 0:1.82:23.64 |
| | $k_{ssw}$ ($s^{-1}$) | 0:5:100 |

**Table S1.** Semisolid MT dictionary parameters (261,888 entries).
*The notation x:y:z denotes a discrete range of values from x to z with a step size of y.

| rNOE dictionary (for training the second network) | | |
|---|---|---|
| Water | $T_1$ (ms) | 1200:100:2300* |
| | $T_2$ (ms) | 40:10:110 |
| MT | $T_1$ (ms) | Fixed to water |
| | $T_2$ (ms) | 0.04 |
| | Chemical shift (ppm) | -2.5 |
| | $f_{ss}$ (%) | 0:1.82:23.64 |
| | $k_{ssw}$ ($s^{-1}$) | 0:5:100 |
| rNOE (-3.5 ppm) | $T_1$ (ms) | Fixed to water |
| | $T_2$ (ms) | 5 |
| | Chemical shift (ppm) | -3.5 |
| | $f_s$ (%) | 0.09:0.14:3.64 |
| | $k_{sw}$ ($s^{-1}$) | 16 |
| rNOE (-1.6 ppm) | $T_1$ (ms) | Fixed to water |
| | $T_2$ (ms) | 0.4 |
| | Chemical shift (ppm) | -1.6 |
| | $f_s$ (%) | 0.075:0.118:3 |
| | $k_{sw}$ ($s^{-1}$) | 16 |

**Table S2.** Dictionary parameters used to simulate rNOE at both -3.5 ppm and -1.6 ppm (a total of 34,673,184 entries).
*The notation x:y:z denotes a discrete range of values from x to z with a step size of y.

(1) Cohen, O.; Huang, S.; McMahon, M. T.; Rosen, M. S.; Farrar, C. T. Rapid and Quantitative Chemical Exchange Saturation Transfer (CEST) Imaging with Magnetic Resonance Fingerprinting (MRF). *Magn. Reson. Med.* **2018**, *80* (6), 2449–2463. https://doi.org/10.1002/mrm.27221.
(2) Shmuely, H.; Rivlin, M.; Perlman, O. Quantitative Multi-Metabolite Imaging of Parkinson's Disease Using AI Boosted Molecular MRI. *Npj Imaging* **2025**, *3* (1), 66. https://doi.org/10.1038/s44303-025-00130-x.
(3) Perlman, O.; Ito, H.; Herz, K.; Shono, N.; Nakashima, H.; Zaiss, M.; Chiocca, E. A.; Cohen, O.; Rosen, M. S.; Farrar, C. T. Quantitative Imaging of Apoptosis Following Oncolytic Virotherapy by Magnetic Resonance Fingerprinting Aided by Deep Learning. *Nat. Biomed. Eng.* **2021**, *6* (5), 648–657. https://doi.org/10.1038/s41551-021-00809-7.
